# Functional autapses form in striatal parvalbumin interneurons but not medium spiny neurons

**DOI:** 10.1101/2022.04.01.486668

**Authors:** Xuan Wang, Zhenfeng Shu, Quansheng He, Xiaowen Zhang, Luozheng Li, Xiaoxue Zhang, Liang Li, Yujie Xiao, Bo Peng, Feifan Guo, Da-Hui Wang, Yousheng Shu

## Abstract

Autapses (or self-synapses) selectively form in specific cell types in many brain regions including the neocortex and the hippocampus, where they provide feedback control over self-spiking activities. Previous morphological studies also found putative autapses in medium spiny neurons (MSNs) of the striatum. However, it remains unclear whether striatal neurons indeed form physiologically functional autapses. We performed whole-cell recordings from striatal neurons in acute mouse brain slices, and identify autaptic neurons by the occurrence of prolonged asynchronous release (AR) of neurotransmitter after high-frequency burst of action potentials (APs) in the same cell. To our surprise, we found no autaptic release in all recorded MSNs after the AP burst, even in the presence of Sr^2+^ that should desynchronize and thus prolong synaptic vesicle release. In sharp contrast, we observed robust autaptic AR events in half of the recorded parvalbumin (PV)-positive neurons. Autaptic responses in PV cells were mediated by GABA_A_ receptors, and the AR strength was dependent on the frequency and the number of APs during the burst. Further simulation results show that autapses regulate burst spiking in PV cells by providing self-inhibition and thus shape network oscillation at certain frequencies. Together, we reveal that, distinct from MSNs, striatal PV neurons form functional autapses, activation of which would regulate self-activities in PV cells, and thereby shape MSN firing and network oscillations.

**Author summary:** Synapses, which usually occur between two neurons, are key structures for signal communication in the nervous system. However, some types of neurons form autapses, where a neuron synapses onto itself. Autaptic transmission provides feedback signal regulating self-spiking activities. Neuronal and network activities in the striatum play critical roles in motor control and other brain functions. Previous studies suggest formation of autapses in striatal principal MSNs, but it remains unclear whether striatal neurons form functional autapses. We performed direct recordings from striatal neurons and examined the occurrence of autaptic transmission in acute brain slices. Surprisingly, we did not detect any autaptic responses in MSNs. A large proportion of striatal PV neurons, however, produced robust autaptic GABA release upon high-frequency stimulation, indicating selective formation of autapses in striatal PV cells. Our computation simulations suggest that autapses provide self-inhibition in PV cells and thereby shape activities in MSNs and striatal network, particularly when PV cells discharge at high frequencies corresponding to a high dopamine state. Together, our findings indicate that PV cells, but not MSNs, in the striatum form physiologically functional autapses. Autapses in PV cells could be essential circuit elements in the striatum and contribute to striatal functions, such as motor control.

## Introduction

Striatum is the largest nucleus in the basal ganglia receiving synaptic inputs from different cortical areas, thalamic nuclei and limbic regions [1]. It plays important roles in various cognitive functions including motor control, emotion processing, learning and memory [2, 3]. In striatum, the most abundant cell type is the projecting GABAergic medium spiny neurons (MSNs), accounting for ~90% of total striatal neurons [4]. Their dendrites are densely covered by spines, a distinct morphological feature of this cell type [5]. MSNs receive information from different brain regions and send output signal to other basal ganglia nuclei, forming the well-known direct and indirect pathways associated with motor control and cognitive functions [6]. In addition, striatum contains various types of interneurons with local axonal arborization regulating MSNs and striatal network activities [1]. Striatal interneurons have distinct morphological and electrophysiological characteristics [7]. Among them, parvalbumin (PV)-expressing interneurons are GABAergic cells with smooth dendrites and a non-adapting high-frequency firing pattern. Although much less abundant (less than 5% of the total) than MSNs, PV cells play key roles in striatal information processing by producing feedforward inhibition onto MSNs [8–11]. Abnormal spiking and synaptic activities in MSNs and PV cells would cause malfunction of the whole basal ganglia network and contribute to the development of brain disorders, such as Parkinson’s disease and Huntington’s disease [12, 13].

Previous studies revealed that GABAergic interneurons in the cortex form massive autaptic connections (known as autapses), i.e. synaptic contacts between the axon of a neuron and its own dendrites or soma [14]. Previous and recent findings also showed that glutamatergic projecting neurons form autaptic connections [15, 16]. Synaptic transmission mediated by autapses generates feedback signal after individual action potentials (APs), providing temporally precise self-control of neuronal spiking activities [16, 17]. In addition to synchronous release (SR) of neurotransmitter release tightly coupled with presynaptic AP generation, delayed asynchronous release (AR) also occurs at autapses [18–20]. Because of this feature of AR at autapses or the conversion of SR to AR with Sr^2+^ [21], it is relatively easy to examine whether a neuron form autaptic connections [18].

Previous morphological findings suggest that striatal MSNs may form autaptic contacts [5, 22]. Moreover, *in vivo* recordings also provided indirect electrophysiological evidence that autapses may exist in MSNs [23]. It remains unclear, however, whether MSNs form physiologically functional autapses. Since PV interneurons in the striatum show similar morphological and electrophysiological properties to those in the cortex, it is of interest to know whether striatal PV cells provide feedback regulation of spiking activity via autapses [19, 20].

With the apparent occurrence of asynchronous neurotransmitter release in physiological condition or in the presence of Sr^2+^ in the bath solution, we have shown abundant autaptic connections in cortical pyramidal cells [16]. In this study, with similar experimental protocols, we examined the occurrence of autaptic AR in both PV cells and MSNs. Surprisingly, we found that PV cells, but not MSNs, form functional autapses. Furthermore, our computational simulations of individual neurons and striatal networks suggest functional roles of PV cell autapses in regulating neuronal activities in both self-activity and network oscillations.

## Results

### MSNs do not form functional autapses

To examine whether MSNs and PV cells form functional autapses, we selectively recorded these cells in coronal slices containing the dorsal striatum of PV-CRE::Ai9 mice (P45-70). PV cells were identified by their expression of tdTomato, while MSNs were identified by their characteristic morphological and electrophysiological features. As reported previously [24], MSNs recorded in our experiments showed a medium-sized cell body, spiny dendrites (Fig 1A and B), and a delayed firing pattern in response to current pulses just above the firing threshold (Fig 1C and D). Meanwhile, consistent with the presence of M-current in MSN [25], a hyperpolarizing current pulse at a near-threshold *V*_m_ level would cause rebound firing (Fig 1C). Our recording showed that the resting *V*_m_ of MSNs was −75.7 ± 1.2 mV and the input resistance was 68.8 ± 9.7 MΩ (n = 22 cells). Consistent with previous studies [26, 27], the AP half-width of MSNs (0.83 ± 0.07 ms, n = 22) was much broader than that of PV cells (0.46 ± 0.02 ms, see below).

**Fig 1.**
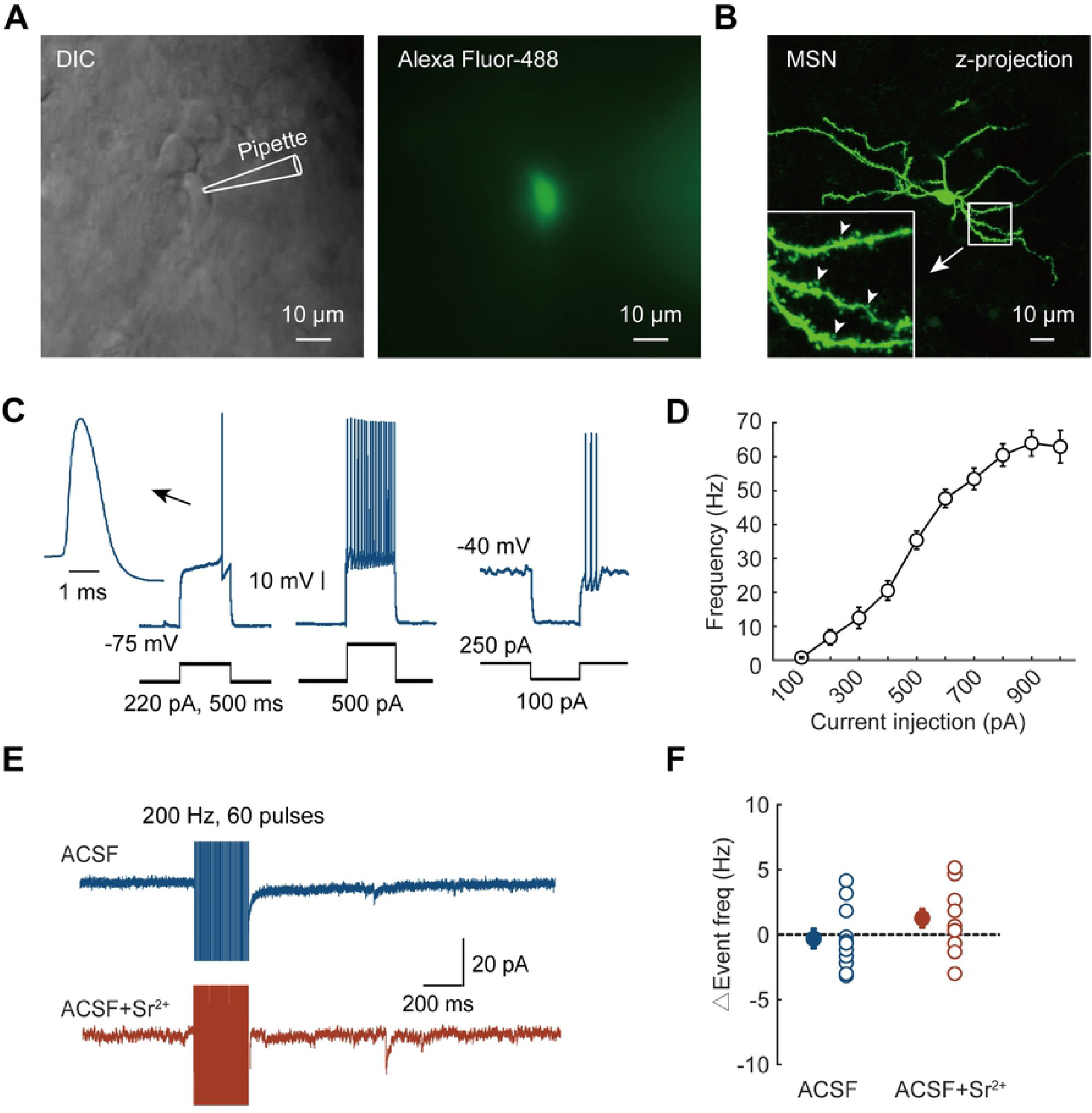
Striatal MSNs do not form functional autapses. (A) DIC (left) and fluorescent images (right) of an MSN loaded with Alexa Fluor-488. (B) A representative image of a recorded MSN with avidin staining. Note the densely distributed spines in the dendrites (inset). Arrowheads indicate some of the spines. (C) Representative traces showing voltage responses to positive step current injections (left and middle) and rebound firing immediately after a hyperpolarizing pulse (right). Inset, expanded action potential. (D) Plot of spike frequency as a function of injected currents (F-I curve) in MSNs. (E) Representative current traces in voltage clamp mode. Note that there was no obvious change in IPSC event number before and after the train stimulation in two conditions (with and without Sr^2+^). (F) Group data showing changes in IPSC event frequency after the train stimulation (see the method section). Note that none of the increments exceed 10 Hz. ns, not significant. Error bars represent s.e.m.

In voltage clamp mode, the membrane potential (*V*_m_) of MSNs were held at −70 mV. With a high-Cl^-^ (~75 mM) internal solution in patch pipettes, inhibitory postsynaptic currents (IPSCs) should be inward at the holding potential (calculated reversal potential: −15 mV). Since fast glutamatergic synaptic events were blocked by kynurenic acid (Kyn, 1.5 mM), all inward synaptic currents should be IPSCs. We stimulated MSNs with trains of brief voltage pulses (50-100 mV, 1 ms in duration, up to 60 pulses) at frequencies from 50 to 200 Hz. As reported previously in neocortical neurons [16, 18], if the recorded neuron form functional autapses and show prolonged AR, or autapses with SR only but in the presence of Sr^2+^, an increase in synaptic events immediately after the train stimulation should be detected. Surprisingly, we found no significant increase in IPSC event number after the train stimulation in both experimental conditions, with or without 5 mM Sr^2+^ in the bath (Fig 1E). With train stimulation of 60 pulses at 200 Hz in the absence of Sr^2+^ (normal ACSF), the increment of IPSC event frequency after the stimulation was −0.30 ± 0.73 Hz (n = 11, Fig 1F). Similar results were obtained in the presence of Sr^2+^ (1.26 ± 0.69 Hz, n = 12). Therefore, distinct from previous morphological observations [5, 22] and indirect electrophysiological evidence [23], our results indicate that striatal MSNs tend not to form functional autapses.

### Autapses form in striatal PV cells

We performed similar experiments in PV cells with tdTomato expression. Close examination of their morphology revealed that PV cells possessed smooth dendrites and dense axon collaterals (Fig 2A and B), similar to previous studies [7, 9, 28]. PV cells had a resting potential of −73.3 ± 1.3 mV (mean ±s.e.m., n = 26 cells) and input resistance of 87.3 ± 9.1 MΩ. They showed unique electrophysiological properties, including non-adaptive fast-spiking pattern (up to 199 ± 18 Hz, n = 26 cells, Fig 2C and D) and short duration of AP waveforms (half-width: 0.46 ± 0.02 ms). Consistent with previous findings, most of the recorded PV cells exhibited stutter firing pattern in response to a series of current steps with increasing current amplitudes (Fig 2C) [28].

**Fig 2.**
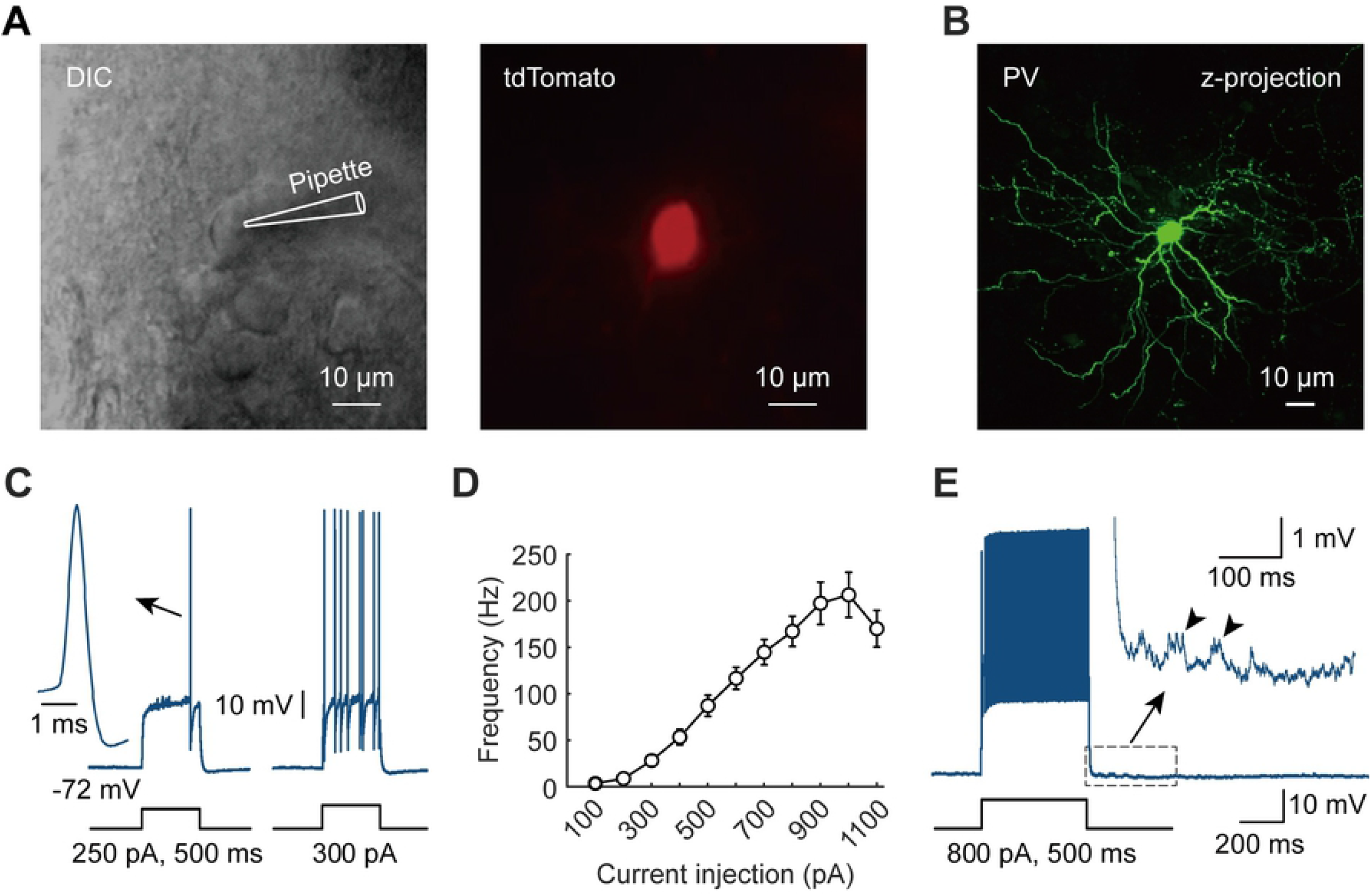
Electrophysiological properties and autaptic responses in PV cells. (A) DIC (left) and tdTomato fluorescent image (right) of a recorded PV cell. (B) A representative image of a recorded PV cell with post hoc staining. (C) Example voltage responses evoked by current pulses. Note the generation of a single AP (left) and the stutter firing pattern (right) evoked by the indicated current steps. Inset, expanded action potential. (D) F-I curve of PV cells. (E) Barrages of autaptic events (arrowheads) occurred immediately after the cessation of AP burst evoked by a strong current pulse. Error bars represent s.e.m.

We frequently observed voltage fluctuations after AP bursts evoked by positive current steps (Fig 2E). These fluctuations contained barrages of depolarizing postsynaptic potentials (PSPs). Since fast glutamatergic synaptic events were blocked by 1.5 mM Kyn, these PSPs should be inversed inhibitory postsynaptic potentials (IPSPs) at the resting *V*_m_. Since PV cells release GABA at their axon terminals and usually hyperpolarize the postsynaptic neurons under physiological conditions, the discharge of PV cells would unlikely drive other neurons to generate APs [19, 20]. Therefore, the PSP barrages occurred immediately after PV cell burst were unlikely caused by polysynaptic events through the network. They actually reflect asynchronous GABA release at autapses of the recorded PV cell, similar to those of fast-spiking cells in mouse and human neocortex [19].

### Autapses are abundant in PV cells and mediated by GABA_A_ receptors

Similarly, in voltage clamp mode, we observed barrages of autaptic currents immediately after the train stimulation in PV cells (Fig 3A). In response to a train of stimulation with 60 APs at 200 Hz, autaptic events reflecting post-train asynchronous release (PT-AR) of GABA lasted for 329 ± 35 ms with a total number of 11.0 ± 1.3 events (n = 55 cells, Fig 3A, see Methods). These autaptic events could be completely blocked by the bath application of GABA_A_ receptor antagonist, picrotoxin (PTX, 50 μM, n = 10, Fig 3B and C), indicating that the autaptic transmission is mediated by GABA_A_ receptors.

**Fig 3.**
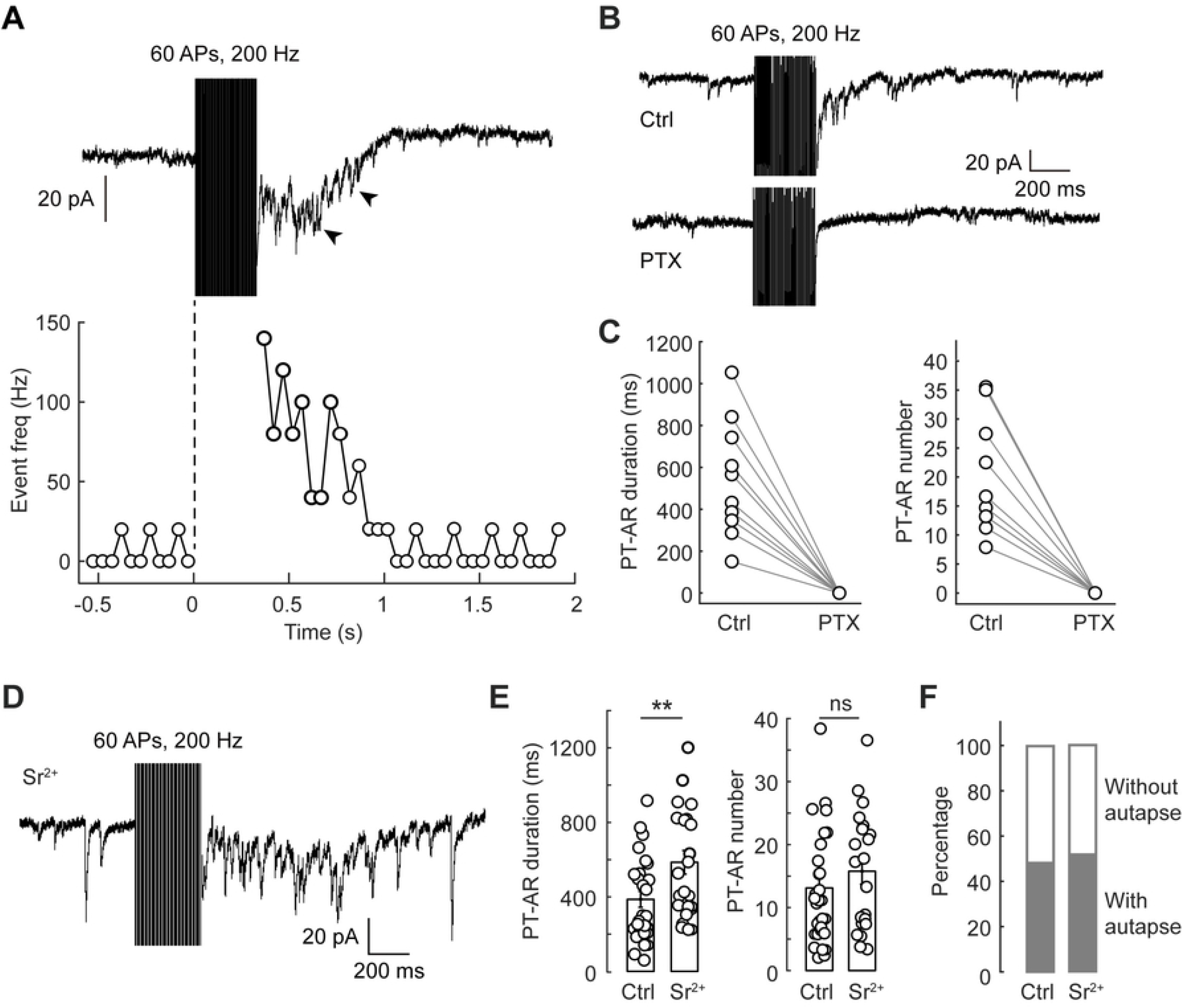
Autaptic transmission is mediated by GABA_A_ receptors and autaptic PV cells are abundant. (A) Top: An example current trace showing the barrages of post-train asynchronous release (PT-AR) events (arrowheads). The PV cell was stimulated with voltage pulses to evoke 60 APs at 200 Hz (holding potential: −70 mV). Bottom: Plot of synaptic event frequency versus time (bin size: 50 ms). The dotted line indicates the onset of AP burst. (B) Example traces before and after the application of 50 μM PTX. (C) Group data showing the effect of PTX on PT-AR duration and event number. (D) An example current trace showing the occurrence of PT-AR in a recorded PV cell bathed with Sr^2+^-containing ACSF. (E) Group data comparing the PT-AR duration and event number in two conditions (Ctrl vs. Sr^2+^). (F) The percentage of autaptic cells in all recorded PV-positive neurons. **, p < 0.01.

Previous studies showed that asynchronous neurotransmitter release occurs selectively in certain types of synapses. For example, AR is much stronger in output synapses of neocortical pyramidal cell onto somatostatin-containing neurons as compared to those onto PV neurons [29]; hippocampal granule cells receive greater AR from CCK cells than that from PV neurons [30]. To exclude the possibility that some autaptic PV cells might have only SR, but no detectable AR, we added SrCl_2_ (5 mM, see Methods) to the bath solution so that autaptic GABA release could be desynchronized [21] (Fig 3D). In our experiments, the presence of Sr^2+^ significantly increased the PT-AR duration from 387 ± 43 to 586 ± 63 ms (control, n = 27; Sr^2+^, n = 21, P = 9.97×10^-3^, two-sample Student’s *t* test). The number of AR events also slightly increased from 13.1 ± 1.7 to 15.8 ± 2.1, but with no significant difference (P = 0.31, Wilcoxon rank sum test, Fig 3E).

We next sought to examine the percentage of PV cells that form autapses. In recorded PV cells, we applied high-frequency stimulations (20-60 APs, 150-200 Hz), with or without Sr^2+^ in ACSF, and monitored the occurrence of post-train autaptic events (see Methods). With this method of AR detection, however, we found that the percentage of PV cells with autaptic AR in Sr^2+^ solution (50.0%, n = 21/42) was similar to that in control condition (without Sr^2+^, 46.6%, n = 61/131, Fig 3F), and also similar to that found in cortical PV neurons [19]. These probabilities should be underestimated because slice preparation reduced the complexity of neuronal dendrites and axons. Together, these results indicate that autapses are abundant in striatal PV cells and autaptic AR occurs in almost every autaptic cell.

### Autaptic AR strength depends on stimulation intensity

Next, we investigated the dependence of autaptic AR strength on the intensity of neuronal activity. We changed the number and the frequency of voltage pulses and monitored the strength of autaptic AR by measuring the PT-AR duration and counting AR events. Consistent with previous findings, the strength of autaptic AR was positively correlated with the number or frequency of the stimuli [18] (Fig 4A and B). When the number of stimulation pulses (200 Hz) increased from 20 to 40 and 60, the average duration of PT-AR increased from 50.6 ± 24.7 to 151 ± 58 and 251 ± 81 ms (ANOVA and post-hoc Tukey’s test, n = 12, P = 0.034), the average number of AR events increased from 2.31 ± 1.28 to 5.33 ± 2.30 and 9.67 ± 3.44 (ANOVA and post-hoc Tukey’s test, n = 12, P = 0.044). Similar dependence of AR strength was observed when we increased the frequency of stimulation pulses. We stimulated the PV cells with a range of frequencies (from 20 to 200 Hz) and found a progressive increase in the PT-AR duration (ANOVA and post-hoc Tukey’s test, n = 15, P = 8.66×10^-7^) and event number (n = 15, P = 5.68×10^-6^, Fig 4C and D). In response to 60 APs at 50 Hz, the average duration and event number of PT-AR were 19.8 ± 9.2 ms and 0.81 ± 0.38, respectively, significantly less than those at 200 Hz (204 ± 41 ms, P = 2.14×10^-4^; 6.50 ± 1.48, P = 5.56×10^-4^, n = 15).

**Fig 4.**
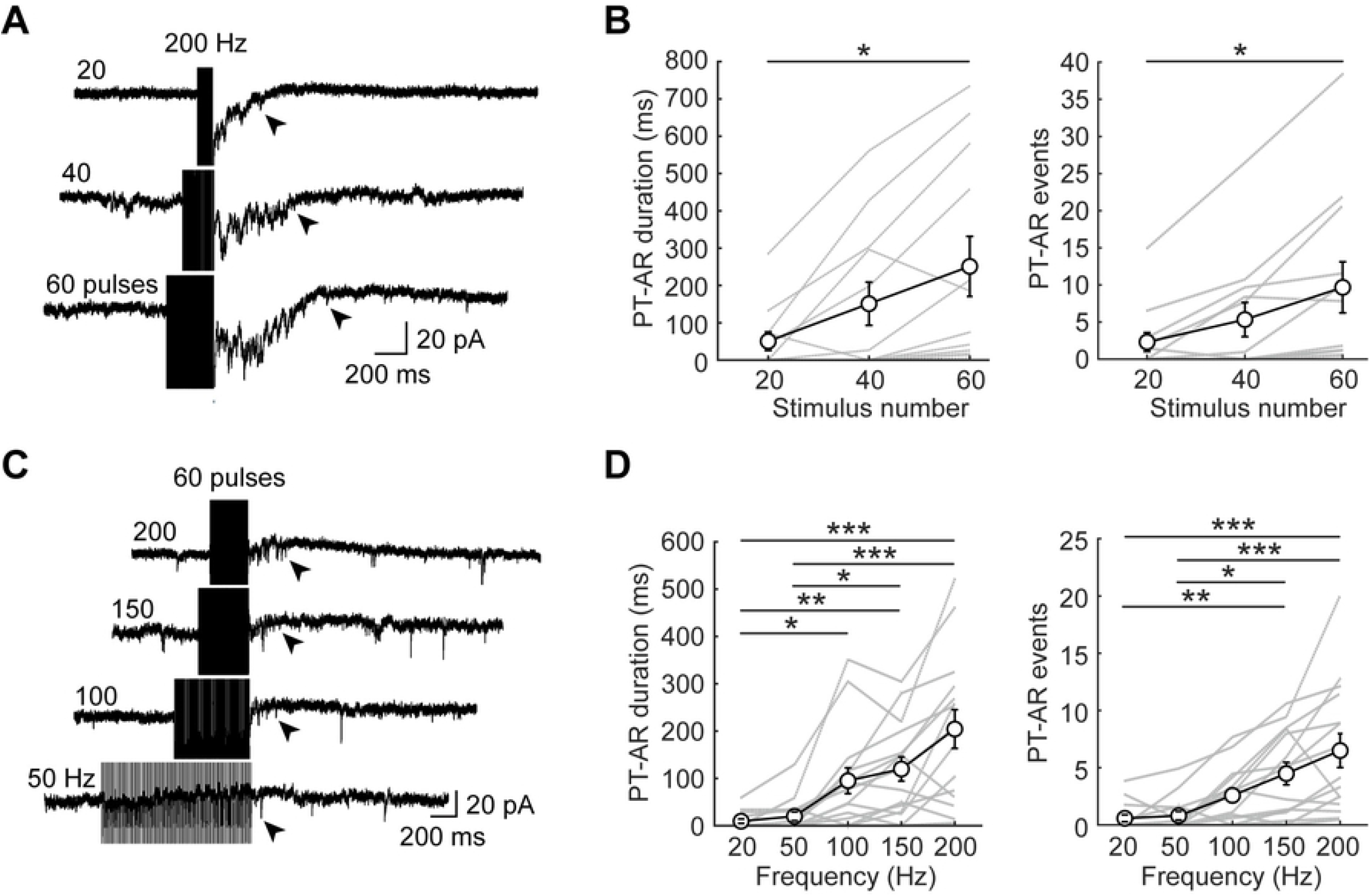
The strength of autaptic AR is dependent on spiking intensity of PV cell. (A) Representative traces showing autaptic currents with different stimulation pulse number (at 200 Hz). Arrowheads indicate the end time of PT-AR. (B) Group data showing the PT-AR duration and event number in response to different number of stimulation pulses at 200 Hz. (C) and (D) Similar as in A and B, but with 60 pulses at different stimulation frequencies. *, p < 0.05; **, p < 0.01; ***, p < 0.001. Error bars represent s.e.m.

### Autapse regulates spiking activity of single PV cell

Next, we attempted to explore the physiological roles of autapses in striatal PV cells. Based on mathematical models of individual PV cells and striatum networks containing not only the PV interneurons but also the principal cells, D1 and D2 MSNs [31], together with models of SR and AR [32, 33], we examined the specific functions of synchronous GABA release or its combination with AR (i.e. SR alone, or SR+AR) in regulating spiking activity of PV cells and MSNs.

We added autapses (with or without AR) to the dendritic compartment of the PV neuron model (see Methods) and compared the differences in firing rate and profile of AP burst (Fig 5A-C). Similar to previous studies [34], our PV cell model also showed two distinct electrophysiological characteristics, stuttering and γ resonance (Fig 5B and C). The strength of autaptic transmission was set to experimental observations (Fig 3A) [19, 33]. Considering that the AR strength was underestimated because some of the autaptic contacts were lost during slicing procedures, we set the standard AR parameters (AR = 1, corresponding to the model parameter τ_AR_ = 150 ms) similar to the strongest AR observed in our experiments, reflecting a condition that the dendrite branches and axon collaterals were relatively more preserved (Fig 5D). In agreement with the experimental findings, the strength of simulated AR (both PT-AR event number and duration) in PV cell model showed dependence on the stimulus number and frequency (Fig 5E and F).

**Fig 5.**
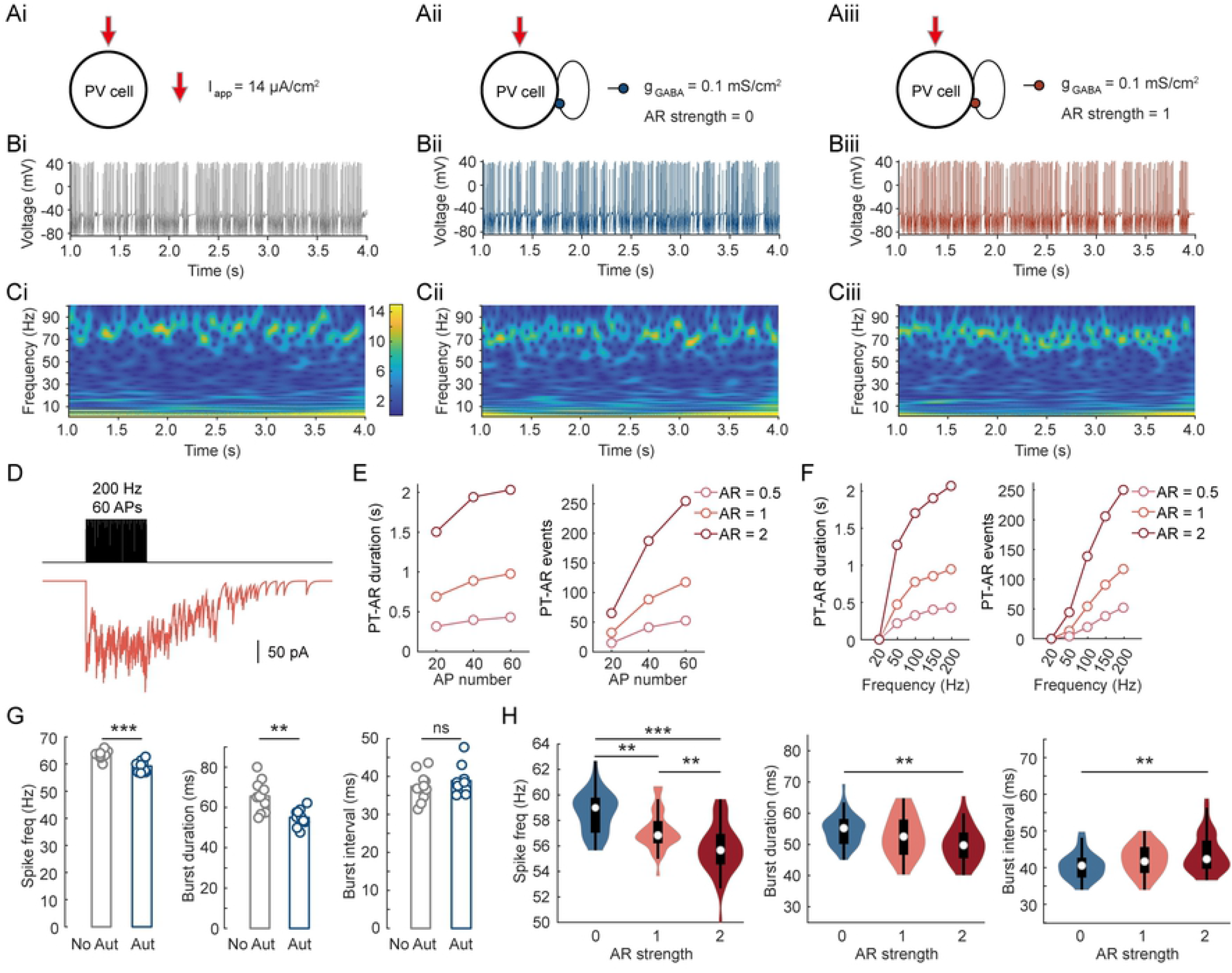
Autapses regulate spiking activity in a single PV cell (simulations). (A) Schematics showing three simulation conditions, PV cell without autapse (No Aut, i), with autapse (Aut, i.e. SR alone, ii) and autaptic AR (SR+AR, iii), in a model of single PV cell. The PV cell received tonic excitation (14 μA/cm^2^) and Poisson noise with a rate of 100 inputs per second (see Methods). (B) Spiking activities of the PV cell model in corresponding conditions shown in A. (C) Spectrograms of the voltage traces in B. (D) An example trace of autaptic currents with AR strength = 1 (i.e. τ_AR_ = 150 ms) when the PV neuron was allowed to discharge 60 APs at 200 Hz. (E) Plots of PT-AR duration and event number as a function of the number of stimuli at 200 Hz (n = 100 trials). Note the increase in duration and event number as the AR strength increased from 0.5 to 1 and 2. (F) Similar as in E, but with 60 APs at different stimulation frequencies. (G) Group data showing the effects of autapses (SR only) on the spiking frequency, burst duration and interval in single PV cell. (H) Group data comparing the firing rate and burst profile with different AR strength. **, p < 0.01; ***, p < 0.001; ns, not significant.

In order to mimic two physiological conditions, i.e. baseline and high dopamine (DA) states, we injected two different background currents (7 and 14 μA/cm^2^) together with the same noise current (Poisson noise) to the modeled PV cell. We then compared the firing patterns with (two conditions: SR only, SR+AR) and without autapses (Fig 5B). At the baseline DA level (S1 Fig), autaptic SR alone decreased the spike frequency and the duration of spike burst, but increased the interval between bursts. However, adding AR showed no further effect on these spiking properties. At the high DA level (Fig 5), we found that SR alone reduced the average firing rate (n = 10 trials, P = 5.75×10^-5^, two-sample Student’s *t* test) and the average duration of bursts (P = 2.00×10^-3^, two-sample Student’s *t* test), but had no significant effect on the burst interval (P = 0.47, Wilcoxon rank sum test, Fig 5G). When we added AR to the autapse (SR+AR), we also observed a slight decrease in the firing rate (n = 40 trials, P = 1.21×10^-9^, ANOVA) and the burst duration (P = 1.31×10^-3^) as the AR strength increased from 0 to 2. Increasing AR strength also prolonged the interval between bursts significantly (P = 7.87×10^-3^, Fig 5H).

### PV cell autapses regulate MSN firing and striatal oscillations

Next, we examined the functional role of autapses in PV cells in the regulation of striatal neuronal and network activity. In a simulated network composed of 50 PV neurons, 100 D1 MSNs and 100 D2 MSNs, we also set the input currents of PV cells to 7 and 14 μA/cm^2^ to mimic the baseline and high DA states, respectively. At the baseline DA level (S2 Fig), PV cells discharged at low frequencies (~9 Hz), adding autaptic SR alone (SR alone, *U_sr_* = 1, *τ_sr_* = 20 ms) exerted marginal effect on PV cell spiking activity and the power at beta and low gamma bands. As expected, adding AR (SR+AR, τ_AR_ = 150 and 300 ms for AR strength 1 and 2, respectively) showed no further effect on both neuronal and network activity (S2 Fig), due to the absence of AR at low firing rates (Fig 4 and 5).

At the high DA level (Fig 6), PV cells discharged around 40 Hz, consistent with that found in movement state [35]. Similar to single PV cell simulations, adding autapses had no significant effect on burst interval (P = 0.070, two-sample Student’s *t* test) but significantly reduced burst duration (P =7.01×10^-9^) and the average spiking frequency of PV cells in the network model (n = 10 trials, P = 1.83×10^-4^ for PV cells, two-sample Student’s *t* test, Fig 6A-G). By contrast, autaptic SR alone increased the activity in both D1 (n = 10 trials, P = 3.59×10^-5^, two-sample Student’s *t* test) and D2 MSNs (n = 10 trials, P = 2.66×10^-4^). At the network level, the power density of certain frequencies in the gamma bands (75-85 Hz) showed a dramatic decrease (n = 10 trials, P = 1.83×10^-4^, Wilcoxon rank sum test), but those of other frequency bands were significantly increased (Fig 6E and G).

**Fig 6.**
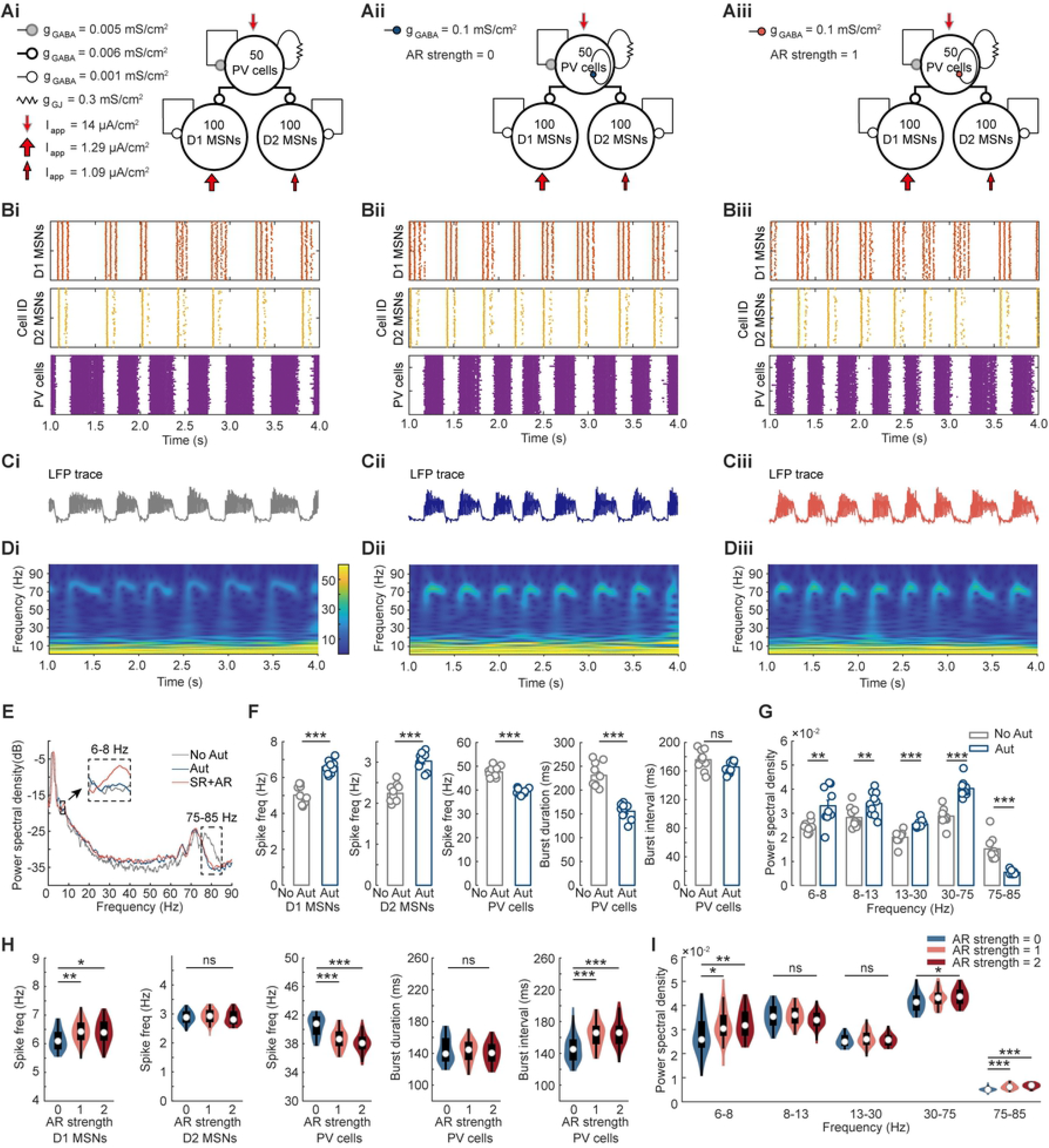
PV cell autapses regulate striatal neuronal and network activities (simulations). (A) Schematics showing three simulation conditions in striatal network model: PV cells without autapse (No Aut, i), with autapse (Aut, SR alone, ii) and autaptic AR (SR+AR, iii). The network model contained 50 PV cells, 100 D1 and 100 D2 MSNs. PV cells received tonic excitation (14 μA/cm^2^) and Poisson noise, while D1 and D2 MSNs received tonic excitation with a strength of 1.29 and 1.09 μA/cm^2^, respectively, corresponding to a high dopamine state. AR strength 1 corresponds to τ_AR_ =150 ms. (B) Raster plots of the three types of striatal neurons in the corresponding conditions shown in A. (C) Example local field potential (LFP) traces in the three conditions. (D) Spectrograms of the LFP traces in C. (E) Mean power spectral analysis of the LFP traces in three conditions. (F) Group data showing the effect of autapses (SR alone) on the firing rate of distinct cell types and the PV cell burst duration and interval. (G) Changes of the power density at LFP frequencies. (H) Group data comparing the firing rate, burst duration and interval with different AR strengths. (I) Changes of the power density at indicated frequencies. *, p < 0.05; **, p < 0.01; ***, p < 0.001; ns, not significant.

In following simulations, we added autaptic AR to PV cells. We found that autaptic AR slightly decreased the firing rate of PV neurons (n = 40, P = 3.18×10^-5^, ANOVA, Fig 6H). In sharp contrast to SR alone, AR had no effect on burst duration (P = 0.563, ANOVA) but slightly increased the burst interval (P = 6.61×10^-5^, ANOVA, Fig 6H). A marginal increase in firing frequency was observed in D1 MSNs, but not D2 MSNs. An increase in AR strength significantly enhanced the power density of theta (6-8 Hz, P = 5.78×10^-3^, ANOVA) and gamma bands (30-75 Hz, P = 0.0133; 75-85 Hz, P = 2.07×10^-8^), but not those of other bands (Fig 6I). Together, these simulations suggest a role of autaptic AR in the regulation of striatal neuronal and network activities.

## Discussion

In this study, we show that autaptic contacts occur in PV interneurons in the striatum. By contrast, no functional autaptic connections were found in the striatal principal cell type, MSNs. Synaptic events mediated by GABA_A_ receptors could be observed after a high-frequency AP burst in PV cells, reflecting asynchronous GABA release at its autapses. We further found that the AR strength is dependent on the frequency and the number of APs. Our simulation results suggest that autapses regulate spiking activities of PV cells by providing self-inhibition. At the network level, activation of PV cell autapses also regulates spiking activities of MSNs and shapes striatal network oscillations.

PV-positive cells can be found in different brain regions, contributing significantly to information processing in a variety of brain circuits [36]. In the neocortex and hippocampus, PV neurons have been confirmed to have functional autapses; similar to certain types of conventional synapses, these autapses possess two modes of neurotransmitter release, synchronous and asynchronous mode [17, 19, 37]. Recordings from acute cortical slices revealed that the percentage of autaptic PV cells in rodent (~85%) is more than that found in human (64.3%) [17, 19]. These percentages could be underestimated because slicing procedures would cut some neurites and thus reduce the complexity of dendritic and axonal branches. In striatum, we found that about half of the recorded PV cells formed autaptic connections in normal physiological condition (i.e. normal bath solution at body temperature). Since adding Sr^2+^ to the bath solution desynchronizes neurotransmitter release and thus delays the occurrence of autaptic events [15, 16, 18], the percentage of autaptic PV cells should be more accurate. However, the percentage with Sr^2+^ was similar to that found in normal bath solution, indicating that AR occurs in physiological conditions.

Do principal MSNs form autapses? Early studies in 1970s reported that some MSN axon collaterals target the soma or proximal dendrites of the same cell in monkey striatum [5]. Later, Park and colleagues performed intracellular recordings in rats and found the occurrence of recurrent inhibition in the recorded MSN when they stimulated the substantial nigra, providing indirect evidence for possible existence of autapses [23]. In cultured MSNs, Shi and Rayport observed autaptic PSPs [22]. In our experiments, we revisited this early question in striatal slices obtained from young adult mice. Surprisingly, we failed to detect any autaptic responses in the recorded MSNs in both normal and Sr^2+^-contained bath solution. Therefore, our results indicate that MSNs do not form functional autapses, distinct from those previous observations. Reasons for the contradictory findings could be as follows. First, the morphologic intersection of axons and dendrites observed in previous studies does not necessarily mean a synaptic structure. Second, *in vivo* observation of recurrent inhibition in MSNs may result from polysynaptic transmission. Last, cultured neurons may form redundant synaptic connections including autapses. Therefore, we believe that MSNs do not form functional autapses. However, it remains to be further examined with electron microscopy whether MSNs form silent autaptic contacts [38].

What are the functional roles of PV autapses? It has long been thought that autapses would regulate the firing pattern of neurons, thus affecting network oscillations [39]. Synchronous GABA release enhances temporal precision of APs in neocortical PV cells [40]. Autapses could promote synchronized firing in neocortical PV cells, allowing them to follow gamma oscillations [41]. Meanwhile, the autaptic AR decreases PV cell spike reliability and desynchronizes the local network [20, 40]. In our simulations, we demonstrate that autaptic SR of GABA shortens the burst duration in PV neurons, allowing earlier rebound firing in MSNs. In contrast, adding autaptic AR prolongs the burst interval of PV cells and thus allows wider time window for MSN to discharge. The activation of MSNs and PV cells will lead to a change of network oscillation. Indeed, the high gamma band power was significantly decreased if we added SR. However, in the presence of AR, both theta and gamma power were significantly increased (Fig 6). Since AR only occur at high frequencies of PV cells, it should only play a role at high DA levels. As expected, adding AR showed no additional effect on neuronal and network activity when PV cells discharge at low frequencies (i.e. baseline DA level). Therefore, we mainly focused on physiological contribution of autapses at states when PV cell discharge at high frequencies.

Feedback inhibition provided by autaptic SR would reduce PV cell spike frequency and shorten its burst duration. Because AR occurs not only during the burst but also after the burst, AR-induced prolonged hyperpolarization would prevent the emergence of next burst in PV cells, thus increasing the burst interval and prolonging MSNs firing. Since PV cells show fast-spiking firing pattern and are able to discharge up to 220 Hz (Fig 2), they contribute largely to high-frequency LFP oscillations. Indeed, we found a decrease in PV cell spike frequency and a shift of power density from high gamma to other bands after adding autaptic SR (Fig 6E-G). Since the frequency of AR events would change progressively after PV cell burst, it should shape LFP oscillations at different bands. We observed increases of gamma band power after introducing AR to the network model. The accumulation of AR events as a whole would cause slow hyperpolarization after PV cell burst, which may contribute to the enhancement of theta band power. Therefore, PV cell autapses shape neuronal activity in both PV cells and MSNs, and regulate network activity via both SR and AR.

Different bands of network activity may play distinct roles in brain functions. The theta band oscillation has been linked to cognitive behavioral states, such as working memory and decision-making [42, 43]. Gamma oscillation is associated with the initiation of movement [44]. Striatal theta oscillation coherence between other brain regions such as hippocampus and amygdala has been shown to facilitate information exchange, and gamma oscillation helps to organize the active neurons in various brain regions [45]. The contribution of PV cell autapses to these bands may play critical roles in proper motor execution and cognition. It has been speculated that dysfunction of interneurons in the striatal network could be an important mechanism for neurological diseases such as Parkinson’s disease [46]. It remains to be examined whether the strength of autaptic connections and their role in the regulation of network oscillations are pathologically altered in different brain disorders.

Together, our results show striatal PV interneurons, but not MSNs, develop functional autapses. Considering the important role of PV cells in the striatal network, we believe that their autapses and the two GABA release modes are fundamental circuit elements and physiological mechanisms, contributing to basal ganglia functions such as motor control and emotion processing. In addition, PV cell autapses could be a key target for the development of new therapies for striatum-related diseases. Our findings also suggest that autaptic connections should be considered when interpreting the function of basal ganglia and building computational models.

## Materials and methods

### Ethical statement

For each experiment, animals of similar age were randomly assigned. The use and care of experimental animals are in line with the guidelines of the Animal Advisory Committee at Fudan University and the State Key Laboratory of Cognitive Neuroscience and Learning, Beijing Normal University.

### Slice preparation

Wild-type C57BL/6J mice and PV-CRE::Ai9 mice (postnatal 45-70 days) were used to prepare striatal slices. Animals were housed with *ad libitum* access to water and food and with a 12-h light/12-h dark cycle. We anesthetized the mouse with sodium pentobarbital (50 mg/kg, intraperitoneal injection), followed by decapitation when there was no sensorimotor reflex. We dissected out the brain and immersed it into 0 °C aerated (95% O_2_ and 5% CO_2_) sucrose-based ACSF (i.e. without NaCl, but with 126 mM sucrose) slicing solution. Tissue blocks were cut coronally with a thickness of 300 μm in this solution using a vibratome (Leica VT1200S). Slices were collected and then incubated in aerated ACSF at 35 °C. The ACSF contained (in mM) 126 NaCl, 2.5 KCl, 2 MgSO_4_, 2 CaCl_2_, 26 NaHCO_3_, 1.25 NaH_2_PO_4_ and 25 dextrose (315 mOsm, pH 7.4). After 40 min of incubation, slices were maintained in the same solution at room temperature until use.

### Electrophysiological recording

Slices were transferred to the recording chamber perfused with aerated ACSF (34 ~ 35 °C) at a rate of 1.2 ml/min. Striatal neurons were visualized using an upright infrared differential interference contrast microscope (BX51WI or BX61WI with two-photon imaging system, Olympus). MSNs were identified by their medium-sized cell bodies and densely distributed dendritic spines, together with their electrophysiological properties (see the main text). PV cells were identified by the expression of tdTomato, together with their fast-spiking firing pattern and narrow AP waveforms. After recording, the recorded neurons were further identified using avidin staining.

Patch pipettes had impedance of 5-8 MΩ when filled with a high Cl^-^ internal solution containing (in mM) 71 KCl, 72 K-gluconate, 2 MgCl_2_, 2 Na_2_ATP, 10 HEPES 0.2 EGTA, and 0.2% biocytin (286 mOsm, pH 7.2). The reversal potential of Cl^-^ was approximately −15 mV. In order to better visualize the morphology of recorded neurons during recording, we added Alexa Fluor-488 (50 μM) to the internal solution. Whole-cell recording was achieved using a MultiClamp 700B amplifier (Molecular Devices). Signals were filtered at 10 kHz and then sampled by Micro 1401 micro3 at 50 kHz using Spike2 software. Firing patterns and membrane properties were examined by step current injections (500 ms in duration, −100 pA to 1,100 pA in amplitude) in current clamp mode. To examine the occurrence of autaptic AR, we stimulated the recorded cell with trains of brief step voltage pulses (10-60 pulses, 0.5-2 ms in pulse duration, 20-200 Hz) in voltage clamp mode. Amplitude of voltage pulses was carefully adjusted to ensure successful generation of APs (i.e. action currents in voltage clamp mode) for each brief pulse.

Kynurenic acid (1.5 mM) was added to the bath solution to block fast glutamatergic transmission (mediated by both NMDA and AMPA receptors). In the Sr^2+^ experiments, we added 5 mM SrCl_2_ to ACSF, but reduced the concentration of both CaCl_2_ and MgSO_4_ to 1 mM. In some experiments, we perfused the slices with 50 μM picrotoxin (PTX, Tocris) to examine whether autaptic responses were mediated by GABA_A_ receptors.

After recordings, slices were fixed with 4% paraformaldehyde (PFA, for more than 12 h) and stained with Alexa Fluor-488 conjugated avidin. The z stack images (0.7 μm between successive images) of individual cells were acquired by a confocal microscope (A1 plus, Nikon) equipped with a 60× objective.

### Computational Models

Neuron models of striatal PV interneurons and MSNs were similar to those previously reported [31]. Synaptic currents (*I_GABA_*) mediated by synchronous and asynchronous GABA release are formulated as in our previous work [33] and that from Volman and colleagues [32]. The model parameters can be found in Table 1.

**Table 1.** Description of the computational models. Models are summarized in panel A and detailed in panels B-G.

| A Striatal Net Work Model Summary |  |
| --- | --- |
| **Populations** | PV cells, D1 MSNs and D2 MSNs |
| **connectivity** | Random convergent connections |
| **Neuron model** | Hodgkin-Huxley model |
| **Transmission modes** | Synchronous release (SR) and asynchronous release (AR) |
| **Input** | A constant current, Poisson noise and Gaussian noise |
| **Measurements** | Spike activity, spike burst and local field potential (LFP) |

| B Populations |  |
| --- | --- |
| Name | Size |
| PV neuron | N = 50 cells |
| D1 MSN | N = 100 cells |
| D2 MSN | N = 100 cells |

| **C** |  |  |  |
| --- | --- | --- | --- |
| **Connectivity** |  |  |  |
| **Name** | **Source** | **Target** | **Pattern** |
| PV-PV<sub>GABA</sub> | PV cell | PV cell | Chance of synaptic connection: 58%<br>$g_{GABA} = 0.005$ mS/cm<sup>2</sup> for high Dopamine (DA) state and $g_{GABA} = 0.1$ mS/cm<sup>2</sup> for baseline DA state |
| PV-PV<sub>GJ</sub> | PV cell dendrite | PV cell dendrite | Chance of gap junction: 33%<br>$g_{GJ} = 0.3$ mS/cm<sup>2</sup> for high DA state and $g_{GJ} = 0.15$ mS/cm<sup>2</sup> for baseline DA state |
| PV<sub>Aut</sub> | PV cell | Dendrite of itself | Chance of connection: 100% for autapse conditions (SR alone and SR + AR); 0 for no autapse conditions<br>$g_{GABA} = 0.1$ mS/cm<sup>2</sup> |
| MSN-MSN | MSN | MSN | Chance of connection: 100%<br>$g_{GABA} = 0.001$ mS/cm<sup>2</sup> |
| PV-MSN | PV cell | MSN | Chance of connection: 37.5%<br>$g_{GABA} = 0.006$ mS/cm<sup>2</sup> |

| **D** |  |
| --- | --- |
| **Neuron Model** |  |
| **Name** | **Function** |
| The dynamics of membrane potential ($V_m$) of each neuron | $c_m \frac{dV}{dt} = -\sum I_{memb} - \sum I_{syn} + I_{inp}$<br>$V$ (membrane voltage) has units of mV.<br>$I$ (Currents) has units of $\mu A/cm^2$.<br>$c_m$ (membrane capacitance) = 1 mF/cm<sup>2</sup><br>$I_{memb}$: intrinsic membrane currents<br>$I_{syn}$: synaptic currents<br>$I_{inp}$: current input |
| The membrane currents | $I_{memb} = \bar{g}(m^n h^k)(V - E_{ion})$<br>$\bar{g}$: a constant maximal conductance<br>$E_{ion}$ is a constant reversal potential<br>The activation ($m$) and inactivation ($h$) gating |
| | variables have $n^{th}$ and $k^{th}$ order kinetics. |
| The dynamics of each gating variable | $\frac{dm}{dt} = \frac{m_{\infty} - m}{\tau_m}$ |
| The steady-state of gating variable | $m_{\infty} = \alpha_m / (\alpha_m + \beta_m)$ |
| The time constant of decay | $\tau_m = 1 / (\alpha_m + \beta_m)$ |
| **PV cell (two compartments)** |  |
| The $V_m$ of somatic ($V$) and dendritic compartment ($V_d$) | $c_m \frac{dV}{dt} = -I_{Na} - I_K - I_L - I_D - I_{syn} + I_{ds}$ $c_m \frac{dV_d}{dt} = -I_{Na} - I_K - I_L - I_D - I_{syn} + I_{inp} + I_{sd}$ |
| Steady state of gating variables of sodium current ($I_{Na}$) | $m_{\infty} = \frac{1}{1 + \exp[-(V + 24)/11.5]}$ $h_{\infty} = \frac{1}{1 + \exp[(V + 58.3)/6.7]}$ $\tau_h = 0.5 + \frac{14}{1 + \exp[(V + 60)/12]}$ |
| Somatic sodium currents parameters | $\bar{g}_{Na} = 112.5 \text{ mS/cm}^2$ $E_{Na} = 50 \text{ mV}$ $n_{Na} = 3$ $k_{Na} = 1$ |
| Steady state of gating variables of potassium current ($I_K$) | $n_{\infty} = \frac{1}{1 + \exp[-(V + 12.4)/6.8]}$ $\tau_n = (0.087 + \frac{11.4}{1 + \exp[(V + 14.6)/8.6]}) (0.087 +$ |
| Somatic potassium current parameters | $\bar{g}_K = 225 \text{ mS/cm}^2$ $E_K = -90 \text{ mV}$ $n_K = 2$ $k_K = 0$ |
| Steady state gating variables of D current ($I_D$) | $a_{\infty} = \frac{1}{1 + \exp[-(V + 50)/20]}$ $b_{\infty} = \frac{1}{1 + \exp[(V + 70)/6]}$ |
| Somatic D current parameters | $\bar{g}_D = 4 \text{ mS/cm}^2$ $E_D = -90 \text{ mV}$ $n_D = 3$ $k_D = 1$ |
| Somatic leak current parameters | $\bar{g}_L = 0.25 \text{ mS/cm}^2$<br>$E_L = -70 \text{ mV}$<br>$n_L = 0$<br>$k_L = 0$ |
| Electrical coupling of somatic and dendritic compartment | $I_{sd} = 0.5(V_{soma} - V_{dend})$<br>$I_{ds} = 0.5(V_{dend} - V_{soma})$<br>$I_{sd}$: current from somatic to dendritic compartment<br>$I_{ds}$: current from dendritic to somatic compartment |
| Dendritic conductance parameters | Conductances in dendritic compartment ($g_{Na}$, $g_K$, $g_D$, $g_L$) are 1/10 the strength of those in somatic compartment |
| **MSNs** |  |
| Membrane potential $V_m$ | $c_m \frac{dV}{dt} = -I_{Na} - I_K - I_L - I_M - I_{syn} + I_{inp}$ |
| The rate functions for sodium current activation ($m$) and inactivation ($h$) variables | $\alpha_m = \frac{0.32(V + 54)}{1 - \exp[-(V + 54)/4]}$<br>$\beta_m = \frac{0.28(V + 27)}{\exp[(V + 27)/5] - 1}$<br>$\alpha_h = 0.128 \exp[-(V + 50)/18]$<br>$\beta_h = \frac{4}{1 + \exp[-(V + 27)/5]}$ |
| Sodium current parameters | $\bar{g}_{Na} = 100 \text{ mS/cm}^2$<br>$E_{Na} = 50 \text{ mV}$<br>$n_{Na} = 3$<br>$k_{Na} = 1$ |
| The rate functions for potassium current of the activation gate | $\alpha_m = \frac{0.032(V + 52)}{1 - \exp[-(V + 52)/5]}$<br>$\beta_m = 0.5 \exp[-(V + 57)/40]$ |
| Potassium current parameters | $\bar{g}_K = 80 \text{ mS/cm}^2$<br>$E_K = -100 \text{ mV}$<br>$n_K = 4$<br>$k_K = 0$ |
| Leak current parameters | $g_L = 0.1 \text{ mS/cm}^2$<br>$E_L = -67 \text{ mV}$<br>$n_L = 0$<br>$k_L = 0$ |
| The rate functions for M-current activation gate | $\alpha_m = \frac{Q_s 10^{-4} (V + 30)}{1 - \exp[-(V + 30)/9]}$ $\beta_m = \frac{Q_s 10^{-4} (V + 30)}{1 - \exp[(V + 30)/9]}$ $Q_s = 3.2094$ |
| M current parameters | $\bar{g}_m = 1.25 \text{ mS/cm}^2$ $E_m = -100 \text{ mV}$ $n_m = 1$ $k_m = 0$ |

| E Synaptic transmission modes |  |
| --- | --- |
| GABA current | $I_{GABA} = g_{GABA} s (V - E)$<br>s: the gating variable for GABAergic synaptic transmission<br>E = -80 mV |
| Gap junction current | $I_{elec} = g_{GJ} (V d_j - V d_i)$ |
| GABA conductance gate variables | $\frac{ds(t)}{dt} = -\frac{s(t)}{\tau_{decay}} + \frac{0.5 \cdot q(t) \cdot (1 - s(t))}{\tau_{rise}}$<br>$\tau_{decay} = 13 \text{ ms}$<br>$\tau_{rise} = 0.25 \text{ ms}$ |
| Total transmitter release | $q(t) = q_{sr}(t) + q_{ar}(t)$ |
| SR amount | $q_{sr}(t) = x_0 n_{sr}(t) \sum_k \delta(t - t_k)$<br>$x_0 = 5$ is the amount of neurotransmitter contained in a vesicle<br>$n_{sr}$: number of released vesicles by SR<br>$t_k$: the arriving moment of the $k$th spike |
| AR amount | $q_{ar}(t) = x_0 n_{ar}(t) \sum_m \delta(t - t_m)$<br>$n_{ar}$: number of released vesicles by AR |
|  | $t_m$: the stochastic release time of AR, in each simulation time bin $dt$, the release probability is $P(t=t_m)=0.01$ |
| SR probability | $\frac{du_{sr}(t)}{dt} = -\frac{u_{sr}(t)}{\tau_{sr}} + U_{sr}[1 - u_{sr}(t)] \sum_m \delta(t - t_m)$<br>$u_{sr}$: SR probability caused by the activation of varied calcium sensors<br>$U_{sr} = 1$ denotes the increments of release probability induced by an action potential via SR<br>$\tau_{sr} = 20$ ms |
| AR probability | $\frac{du_{ar}(t)}{dt} = -\frac{u_{ar}(t)}{\tau_{ar}} + U_{ar}[1 - u_{ar}(t)] \frac{(Ca(t) - Ca_{baseline})^4}{(Ca(t) - Ca_{baseline})^4 + 1}$<br>$u_{ar}$: AR probability caused by the activation of varied calcium sensors<br>$Ca(t)$: residual calcium in presynaptic terminal at a given time $t$<br>$U_{ar} = 0.4$ denotes the increments of release probability induced by an action potential via AR<br>$\tau_{ar} = 150$ ms for AR strength, condition 1 (SR+AR)<br>$\tau_{ar} = 0.01$ ms for AR strength, condition 0 (Aut, i.e. SR alone)<br>$K_a = 0.8$ is the calcium affinity of AR machinery<br>$Ca_{baseline} = 0.05$ is the baseline intracellular calcium level |
| Available vesicle numbers | $dN(t) = r(t) - n_{ar}(t) \sum_m \delta(t - t_m)dt - n_{sr}(t) \sum_k \delta(t - t_k)dt$ |
| Synaptic vesicle replenishment | $r(t) \sim B(N_F - N(t), dt/\tau_d)$<br>$N_F = 10$ is the total number of vesicles<br>$\tau_d = 15$ is the time constant of vesicle replenishment |
| SR vesicles | $n_{sr}(t) \sim B(N(t), u_{sr}(t_+))$<br>$t_+$: the moment of just after the arrival of an action potential. |
| AR vesicles | $n_{ar}(t) \sim B(N(t), u_{ar}(t))$ |
| Residual calcium level | $\frac{dCa(t)}{dt} = -\beta \frac{Ca^2}{Ca^2 + K_p^2} + \gamma \log \frac{Ca_{outside}}{Ca} \sum_m \delta(t - t_m) + \beta \frac{Ca_{outside}}{Ca}$<br>$Ca_{outside} = 0.002$ is the extracellular calcium concentration<br>$\beta = 0.02$ is the maximal rate with which residual calcium is cleared from synaptic terminal by active pumps<br>$K_p = 0.8$ is the affinity of a pump for calcium<br>$\gamma = 0.001$ controls the amount of per-spike increase in residual synaptic calcium |

| F Input |  |
| --- | --- |
| Name | $I_{inp}$ |
| PV cell | Tonic current input $I_{app} = 14 \mu\text{A}/\text{cm}^2$ for high DA state, and $I_{app} = 7 \mu\text{A}/\text{cm}^2$ for baseline DA state |
|  | Poisson input: 50 uncorrelative homogeneous Poisson processes firing at 2 Hz with synaptic conductance fixed at 0.3 nS |
| MSN | Tonic current input $I_{app} = 1.29 \mu\text{A}/\text{cm}^2$ for D1 MSNs and $I_{app} = 1.09 \mu\text{A}/\text{cm}^2$ for D2 MSNs to model the high DA state, and $I_{app} = 1.19 \mu\text{A}/\text{cm}^2$ for D1 and D2 MSNs at baseline DA state |
|  | Gaussian noise has a mean 0, standard deviation 1 |

| G Measurements |  |
| --- | --- |
| Spike activity | Raster plots and spiking rates |
| Burst | any occurrence of at least two spikes with an inter-spike interval (ISI) less than 16 ms [47] |
| LFP | Sum of all synaptic currents in all cells. To eliminate transients due to initial conditions, LFP is evaluated 1 s after the simulation onset. |

For a single PV neuron, we reduced the number of PV neurons to 1 and focused only on its spiking activity and membrane potential *V*_m_. We simulated the absence of autapses (No Aut) by adjusting the chance of autaptic connection to 0. For autaptic PV cells, we simulated conditions with different strength of AR by adjusting the parameter *τ_ar_*. To examine the contribution of autapses to network oscillation, we normalized (z-score) the simulated LFP before plotting the power spectrum. All simulations were run on MATLAB software (version R2021a). All differential equations were integrated using a fourth-order Runge-Kutta algorithm with time step 0.01 ms.

### Data Analysis

Spike2, MiniAnalysis and MATLAB software were used for data analysis. All measurements were taken from different cells. Unless otherwise stated, data presented in the main text was mean ± s.e.m. The error bars in figures were also s.e.m.

The frequency of spontaneous IPSC events was obtained from 2-s baseline current just before the stimulation onset, and considered as the baseline frequency. For a particular neuron, if the frequency of IPSC events within 300 ms after the train stimulation (60 pulses at 200 Hz) is 10 Hz higher than that of the baseline, it would be considered as an autaptic neuron with asynchronous GABA release. The PT-AR instantaneous frequency was also calculated. The termination time of PT-AR IPSC barrage is the time of the last IPSC event before the AR frequency reaches the baseline frequency. The PT-AR duration is the time period between the cessation of the train stimulation and the end of the AR barrage.

For two independent observations with normal distribution (P > 0.05, Shapiro Wilk’s test), we used two-sample Student’s *t* test. Non-normal data were compared with Wilcoxon rank sum test. Kruskal-Wallis test for analysis of variance (ANOVA), and Tukey’s test for post-hoc analysis, were used for comparisons of multiple groups. Datasets were considered to be significantly different if P < 0.05.

## Data availability statement

All relevant data are within the manuscript and the Supporting Information files. Raw electrophysiological traces are available from the corresponding authors upon request. Computation codes for simulations are available in a GitHub repository at https://github.com/xuanwangbnu/Aut-A-S. DOI: 10.5281/zenodo.6397521.

## Acknowledgments

We are grateful to Drs. Suixin Deng and Junlong Li for their help in data analysis.

## Supporting information

**S1 Fig. Autapses regulate spiking activity in single PV cell (baseline DA state).**

(A) Schematics showing three simulation conditions, PV cell without autapse (No Aut, i), with autapse (Aut, i.e. SR alone, ii) and autaptic AR (SR+AR, iii), in a model of single PV cell. The PV cell received tonic excitation (7 μA/cm^2^) and Poisson noise. (B) Spiking activities of the PV cell model in corresponding conditions shown in A. (C) Spectrograms of the voltage traces in B. (D) Group data showing the effects of autapses on the spiking frequency, burst duration and interval in single PV cell. (E) Group data comparing the firing rate and burst profile with different AR strength. *, p < 0.05;**, p < 0.01; ***, p < 0.001; ns, not significant.

**S2 Fig. PV cell autapses regulate striatal neuronal and network activities (baseline DA state).**

(A) Schematics showing three simulation conditions in striatal network model: PV cells without autapse (No Aut, i), with autapse (Aut, SR alone, ii) and autaptic AR (SR+AR, iii). The network model contained 50 PV cells, 100 D1 and 100 D2 MSNs. PV cells received tonic excitation (7 μA/cm^2^) and Poisson noise, while D1 and D2 MSNs received tonic excitation with a strength of 1.19 μA/cm^2^, corresponding to a baseline DA state. AR strength 1 corresponds to τ_AR_ = 150 ms. (B) Raster plots of the three types of striatal neurons in the corresponding conditions shown in A. (C) Example local field potential (LFP) traces in the three conditions. (D) Spectrograms of the LFP traces in C. (E) Mean power spectral density of the LFP traces in three conditions. (F) Group data showing the effect of autapses (SR alone) on the firing rate of distinct cell types and the PV cell burst duration and interval. (G) Changes of the power density at LFP frequencies. (H) Group data comparing the firing rate, burst duration and interval in PV cells with different AR strengths. (I) Changes of the power density at indicated frequencies. *, p < 0.05; **, p < 0.01; ns, not significant.

## References

1. Burke DA, Rotstein HG, Alvarez VA. Striatal Local Circuitry: A New Framework for Lateral Inhibition. Neuron. 2017;96(2):267–84.

2. Vergara R, Rick C, Hernandez-Lopez S, Laville JA, Guzman JN, Galarraga E, et al. Spontaneous voltage oscillations in striatal projection neurons in a rat corticostriatal slice. J Physiol. 2003;553(Pt 1):169–82. doi: 10.1113/jphysiol.2003.050799.

3. Miguelez C, Morera-Herreras T, Torrecilla M, Ruiz-Ortega JA, Ugedo L. Interaction between the 5-HT system and the basal ganglia: functional implication and therapeutic perspective in Parkinson’s disease. Front Neural Circuits. 2014;8:21. doi: 10.3389/fncir.2014.00021.

4. Ponzi A, Barton SJ, Bunner KD, Rangel-Barajas C, Zhang ES, Miller BR, et al. Striatal network modeling in Huntington’s Disease. PLoS Comput Biol. 2020;16(4):e1007648. doi: 10.1371/journal.pcbi.1007648.

5. DiFiglia M, Pasik P, Pasik T. A Golgi study of neuronal types in the neostriatum of monkeys. Brain Res. 1976;114(2):245–56.

6. Calabresi P, Picconi B, Tozzi A, Ghiglieri V, Di Filippo M. Direct and indirect pathways of basal ganglia: a critical reappraisal. Nat Neurosci. 2014;17(8):1022–30. doi: 10.1038/nn.3743.

7. Silberberg G, Planert H. Optogenetic Dissection of the Striatal Microcircuitry. In: Korngreen A, editor. Advanced Patch-Clamp Analysis for Neuroscientists. New York, NY: Springer New York; 2016. p. 151–70.

8. Blomeley CP, Bracci E. Serotonin excites fast-spiking interneurons in the striatum. Eur J Neurosci. 2009;29(8):1604–14. doi: 10.1111/j.1460-9568.2009.06725.x.

9. Orduz D, Bischop DP, Schwaller B, Schiffmann SN, Gall D. Parvalbumin tunes spike-timing and efferent short-term plasticity in striatal fast spiking interneurons. J Physiol. 2013;591(13):3215–32. doi: 10.1113/jphysiol.2012.250795.

10. Owen SF, Berke JD, Kreitzer AC. Fast-Spiking Interneurons Supply Feedforward Control of Bursting, Calcium, and Plasticity for Efficient Learning. Cell. 2018;172(4):683–95.e15. doi: 10.1016/j.cell.2018.01.005.

11. Johansson Y, Silberberg G. The Functional Organization of Cortical and Thalamic Inputs onto Five Types of Striatal Neurons Is Determined by Source and Target Cell Identities. Cell Rep. 2020;30(4):1178–94.e3. doi: 10.1016/j.celrep.2019.12.095.

12. Mallet N, Ballion B, Le Moine C, Gonon F. Cortical inputs and GABA interneurons imbalance projection neurons in the striatum of parkinsonian rats. J Neurosci. 2006;26(14):3875–84. doi: 10.1523/Jneurosci.4439-05.2006.

13. Cepeda C, Galvan L, Holley SM, Rao SP, Andre VM, Botelho EP, et al. Multiple sources of striatal inhibition are differentially affected in Huntington’s disease mouse models. J Neurosci. 2013;33(17):7393–406. doi: 10.1523/Jneurosci.2137-12.2013.

14. Bekkers JM. Neurophysiology: Are autapses prodigal synapses? Curr Biol. 1998;8(2):R52–R5.

15. Van Der Loos H, Glaser EM. Autapses in neocortex cerebri: synapses between a pyramidal cell’s axon and its own dendrites. Brain Res. 1972;48:355–60.

16. Yin L, Zheng R, Ke W, He Q, Zhang Y, Li J, et al. Autapses enhance bursting and coincidence detection in neocortical pyramidal cells. Nat Commun. 2018;9(1):4890. doi: 10.1038/s41467-018-07317-4.

17. Bacci A, Huguenard JR, Prince DA. Functional autaptic neurotransmission in fast-spiking interneurons: A novel form of feedback inhibition in the neocortex. J Neurosci. 2003;23(3):859–66.

18. Li J, Deng S, He Q, Ke W, Shu Y. Asynchronous Glutamate Release at Autapses Regulates Spike Reliability and Precision in Mouse Neocortical Pyramidal Cells. Cereb Cortex. 2021;31(4):2278–90. doi: 10.1093/cercor/bhaa361.

19. Jiang M, Zhu J, Liu Y, Yang M, Tian C, Jiang S, et al. Enhancement of asynchronous release from fast-spiking interneuron in human and rat epileptic neocortex. PLoS Biol. 2012;10(5):e1001324. doi: 10.1371/journal.pbio.1001324.

20. Manseau F, Marinelli S, Mendez P, Schwaller B, Prince DA, Huguenard JR, et al. Desynchronization of Neocortical Networks by Asynchronous Release of GABA at Autaptic and Synaptic Contacts from Fast-Spiking Interneurons. PLoS Biol. 2010;8(9):e1000492. doi: 10.1371/journal.pbio.1000492.

21. Bekkers JM, Clements JD. Quantal amplitude and quantal variance of strontium-induced asynchronous EPSCs in rat dentate granule neurons. J Physiol. 1999;516 (Pt 1):227–48. doi: 10.1111/j.1469-7793.1999.227aa.x.

22. Shi W, Rayport S. GABA Synapses Formed in vitro by Local Axon Collaterals of Nucleus Accumbens Neurons. J Neurosci. 1994;14(7):4548–60.

23. Park MR, Lighthall JW, Kitai ST. Recurrent inhibition in the rat neostriatum. Brain Res. 1980;194(2):359–69. doi: 10.1016/0006-8993(80)91217-2.

24. Wilson CJ, Groves PM. Fine structure and synaptic connections of the common spiny neuron of the rat neostriatum: A study employing intracellular injection of horseradish peroxidase. J Comp Neurol. 1980;194(3):599–615. doi: 10.1002/cne.901940308.

25. McCarthy MM, Moore-Kochlacs C, Gu X, Boyden ES, Han X, Kopell N. Striatal origin of the pathologic beta oscillations in Parkinson’s disease. P Natl Acad Sci USA. 2011;108(28):11620–5. doi: 10.1073/pnas.1107748108.

26. Venance L, Glowinski J. Heterogeneity of spike frequency adaptation among medium spiny neurones from the rat striatum. Neuroscience. 2003;122(1):77–92. doi: 10.1016/S0306-4522(03)00553-0.

27. Cepeda C, Andre VM, Yamazaki I, Wu N, Kleiman-Weiner M, Levine MS. Differential electrophysiological properties of dopamine D1 and D2 receptor-containing striatal medium-sized spiny neurons. Eur J Neurosci. 2008;27(3):671–82. doi: 10.1111/j.1460-9568.2008.06038.x.

28. Kawaguchi Y. Physiological, morphological, and histochemical characterization of three classes of interneurons in rat neostriatum. J Neurosci. 1993;13(11):4908–23.

29. Deng S, Li J, He Q, Zhang X, Zhu J, Li L, et al. Regulation of Recurrent Inhibition by Asynchronous Glutamate Release in Neocortex. Neuron. 2020;105(3):522–33.e4. doi: 10.1016/j.neuron.2019.10.038.

30. Hefft S, Jonas P. Asynchronous GABA release generates long-lasting inhibition at a hippocampal interneuron-principal neuron synapse. Nat Neurosci. 2005;8(10):1319–28. doi: 10.1038/nn1542.

31. Chartove JAK, McCarthy MM, Pittman-Polletta BR, Kopell NJ. A biophysical model of striatal microcircuits suggests gamma and beta oscillations interleaved at delta/theta frequencies mediate periodicity in motor control. PLoS Comput Biol. 2020;16(2):e1007300.

32. Volman V, Behrens MM, Sejnowski TJ. Downregulation of parvalbumin at cortical GABA synapses reduces network gamma oscillatory activity. J Neurosci. 2011;31(49):18137–48. doi: 10.1523/Jneurosci.3041-11.2011.

33. Wang T, Yin LP, Zou XL, Shu YS, Rasch MJ, Wu S. A Phenomenological Synapse Model for Asynchronous Neurotransmitter Release. Front Comput Neurosc. 2016;9. doi: 10.3389/fncom.2015.00153.

34. Sciamanna G, Wilson CJ. The ionic mechanism of gamma resonance in rat striatal fast-spiking neurons. J Neurophysiol. 2011;106(6):2936–49. doi: 10.1152/jn.00280.2011.

35. Chen H, Lei H, Xu Q. Neuronal activity pattern defects in the striatum in awake mouse model of Parkinson’s disease. Behav Brain Res. 2018;341:135–45. doi: 10.1016/j.bbr.2017.12.018.

36. Hu H, Gan J, Jonas P. Interneurons. Fast-spiking, parvalbumin+ GABAergic interneurons: from cellular design to microcircuit function. Science. 2014;345(6196):1255263.

37. Nahar L, Delacroix BM, Nam HW. The Role of Parvalbumin Interneurons in Neurotransmitter Balance and Neurological Disease. Front Psychiatry. 2021;12:679960. doi: 10.3389/fpsyt.2021.679960.

38. Kerchner GA, Nicoll RA. Silent synapses and the emergence of a postsynaptic mechanism for LTP. Nat Rev Neurosci. 2008;9(11):813–25. doi: 10.1038/nrn2501.

39. Herrmann CS, Klaus A. Autapse turns neuron into oscillator. Int J Bifurcat Chaos. 2004;14(02):623–33. doi: 10.1142/s0218127404009338.

40. Bacci A, Huguenard JR. Enhancement of spike-timing precision by autaptic transmission in neocortical inhibitory interneurons. Neuron. 2006;49(1):119–30. doi: 10.1016/j.neuron.2005.12.014.

41. Deleuze C, Bhumbra GS, Pazienti A, Lourenço J, Mailhes C, Aguirre A, et al. Strong preference for autaptic self-connectivity of neocortical PV interneurons facilitates their tuning to γ-oscillations. PLoS Biol. 2019;17(9):e3000419–e. doi: 10.1371/journal.pbio.3000419.

42. Gui D-Y, Yu T, Hu Z, Yan J, Li X. Dissociable functional activities of cortical theta and beta oscillations in the lateral prefrontal cortex during intertemporal choice. Sci Rep. 2018;8(1):11233. doi: 10.1038/s41598-018-21150-1.

43. Petersen PC, Buzsaki G. Cooling of Medial Septum Reveals Theta Phase Lag Coordination of Hippocampal Cell Assemblies. Neuron. 2020;107(4):731–44.e3. doi: 10.1016/j.neuron.2020.05.023.

44. Masimore B, Schmitzer-Torbert NC, Kakalios J, David Redish A. Transient striatal γ local field potentials signal movement initiation in rats. NeuroReport. 2005;16(18):2021–4.

45. Berke JD. Fast oscillations in cortical-striatal networks switch frequency following rewarding events and stimulant drugs. Eur J Neurosci. 2009;30(5):848–59. doi: 10.1111/j.1460-9568.2009.06843.x.

46. Berke JD, Okatan M, Skurski J, Eichenbaum HB. Oscillatory Entrainment of Striatal Neurons in Freely Moving Rats. Neuron. 2004;43(6):883–96.

47. Payeur A, Guerguiev J, Zenke F, Richards BA, Naud R. Burst-dependent synaptic plasticity can coordinate learning in hierarchical circuits. Nat Neurosci. 2021;24(7):1010–9.

